# Epigenomic landscape of the developing human rhombic lip reveals gene regulatory network and non-coding loci of developmental, evolutionary, and disease relevance

**DOI:** 10.1101/2025.10.30.685586

**Authors:** Xinghan Sun, Soumya Menon, Paul Wambo, Ilinca Lungu, Birth Defects Research Laboratory, Kimberly A. Aldinger, Shraddha Pai

## Abstract

The cerebellar rhombic lip neural progenitor niche of the prenatal hindbrain is an anatomical structure critical for cerebellar glutamatergic neurogenesis. Humans have elaborated the rhombic lip niche to include a rhombic lip subventricular zone (RL-SVZ) not seen in mice or macaques. Although developmental disruptions of this progenitor zone can cause cerebellar growth abnormalities – from malformations to tumors – the gene regulatory networks underpinning this unique progenitor niche are unknown. Here we provide a predicted gene regulatory network for the human cerebellar rhombic lip, inferred from epigenomic maps of the developing human cerebellum. We generated DNA methylomes of neuroanatomically-dissected mid-gestation human rhombic lip ventricular zone and RL-SVZ (N=9 samples; 15-16 post conception weeks) using low-input Enzymatic MethylSeq. We also mapped histone modifications marking active promoters and enhancers in the whole mid-gestation human fetal cerebellum (N=6 samples; 14 and 18 post-conception weeks). Integrating these data, we identified 9,855 differentially-methylated regions (DMR) which converge on binding sites of over three hundred transcription factors, including master regulators of rhombic lip neurogenesis, ATOH1, NEUROD1, and NEUROD2. DMRs hypomethylated in the RL-SVZ are enriched in active enhancers and in human accelerated regions, and are depleted in active promoters. We inferred 81,844 transcription factor-enhancer-gene links, covering 41 transcription factors active in the rhombic lip, and 4,610 target genes that include drivers of cerebellar neurogenesis and pediatric hindbrain cancer. Twenty-five DMRs overlap human accelerated regions located near genes associated with intellectual disability, autism spectrum disorders, and neurological deficits. DMRs are also statistically enriched in copy number aberrations in medulloblastoma, a malignant pediatric hindbrain cancer with subtypes hypothesized to originate in the rhombic lip. Close to one-quarter of the DMRs overlap known copy number aberrations in medulloblastoma, nominating potential enhancer and promoter elements impacted by these genomic aberrations. Collectively, our data provide a rich resource to start decoding the functional impact of non-coding variation on gene regulation in the developing cerebellum and on genomic dysregulation in diseases of cerebellar growth.

## Introduction

While the developing human cerebellum initially shares similarity with non-human primates and mice, important differences emerge in the first trimester; one such difference arises in the rhombic lip neurogenic niche^1–3^. This progenitor zone generates all glutamatergic neurons in the cerebellum, including granule neuron progenitors which produce granule neurons, which comprise over half of all neurons in the adult human brain^4^, and unipolar brush cells. The human cerebellar rhombic lip has two anatomical subcompartments. The first is the SOX2^+^ KI67^+^ rhombic lip ventricular zone. The second is the SOX2^-^ KI67^+^ rhombic lip subventricular zone, which to date has only been seen in the developing human brain, but not mice or macaques^3,5,6^. Developmental dysregulation of the rhombic lip is hypothesized to cause some types of childhood hindbrain cancers such as Group 3 and 4 medulloblastoma^5,7^, as well as cerebellar structural birth defects, such as Dandy-Walker malformation^2^. The rhombic lip is therefore an anatomical structure relevant to cerebellar development, growth disorders, and potentially to human cerebellar expansion across primate evolution.^3,6^

Our goal is to identify gene regulatory networks - enhancers, transcription factors, and target genes - of the rhombic lip ventricular and subventricular zones. Elucidating this network will nominate master regulators of human cerebellar neurogenesis and evolutionary expansion, and identify gene expression programs that may be unique to the human-enriched rhombic lip subventricular zone (RL-SVZ). Annotating the epigenomic landscape of the developing rhombic lip will also allow inference of the impact of non-coding genetic variation in cerebellar disorders, on the transcriptome, and ultimately, cellular phenotype.

The cerebellar rhombic lip is a small region that can be readily identified based on its neuroanatomical location and cellular density, and readily captured using tissue microdissection combined with low-input genome profiling technologies^3^. Here we profiled these compartments by combining laser-capture microdissection of the rhombic lip ventricular and subventricular zones, with Enzymatic MethylSeq (EMseq^8^). EMseq is a sensitive enzyme-based DNA methylation assay that does not damage the tissue as the traditional bisulfite sequencing assay does, and it excels at producing unbiased DNA methylome coverage with low starting input amounts. To nominate *cis* regulatory DNA elements active in the rhombic lip, we integrated the rhombic lip DNA methylomes with histone maps of the whole human fetal cerebellum. We used ChIP-seq to map peaks of H3K27ac - which marks active enhancers - H3K4me3 - which marks active promoters - and RNA Polymerase II, using mid-gestation whole human fetal cerebellum as input. While more predictive of active *cis* regulatory DNA elements^9–11^ than DNA methylation, these assays require orders of magnitude more input DNA, making them currently infeasible to map small tissue compartments such as the rhombic lip. To infer target genes of regions demonstrating changes in DNA methylation in the differentiating rhombic lip, we integrated our epigenomic maps with predictions of enhancer-target gene associations^9,10^, and long-range chromatin interaction maps from a single-cell epigenomic map of the first trimester human brain^12^. We identified 1,400 genomic loci that overlap putative active enhancers, and 1,384 loci that are predicted to regulate 4,604 genes. These genes include 29 neurodevelopment-associated genes and 60 genes known to be recurrently mutated or overexpressed in medulloblastoma. Roughly one-quarter of all epigenetically dynamic loci overlap structural variants in Group 3 and 4 medulloblastoma. Collectively the novel fetal epigenomic maps and target gene predictions represent a data-rich starting point to infer gene regulatory networks driving cerebellar neurodevelopment and ascertain the transcriptomic impact of noncoding variation in diseases affecting cerebellar growth.

## Methods

### Neuroanatomical isolation of mid-gestation rhombic lip cell compartments and generation of DNA methylomes

This project was approved by the Research Ethic Board at the University of Toronto (RIS Human Protocol #42524). We received frozen mid-gestation human hindbrains from the University of Maryland Brain and Tissue Bank (all males; mean age = 16 post-conception weeks, Supplementary Table 1). Tissue cryosectioning and immunohistochemistry was performed by the University Health Network Pathology Research Program Laboratory. Brain tissue was cryosectioned on the mid-sagittal plane and stained for cresyl violet, SOX2 (CST Sox2 (D6D9) XP Rabbit mAb #3579) and KI67 (Dako #M7240 Mouse monoclonal antibody). The rhombic lip was identified at the edge of the fourth ventricle by its size and prominent KI67+ staining (**Fig. 1a**). Selected sections were stained with CD34 (Dako #M7165 Monoclonal Mouse Anti-Human) and GFAP (Dako #Z334) to identify the vascular bed that separates the rhombic lip ventricular zone and subventricular zone (Supplementary Figure 1). Using immunohistochemistry, we identified the rhombic lip ventricular (SOX2^+^ KI67^+^) and subventricular (SOX2^-^ KI67^+^) zone in each sample (**Fig. 1a**). We used laser capture microdissection (LCM) to isolate tissue separately from the rhombic lip ventricular zone and subventricular zone. LCM was performed by the Ontario Institute for Cancer Research Tissue Portal using the Leica LMD6 instrument; cresyl violet was used to visualize the tissue. We extracted nucleic acid from each tissue compartment, and generated DNA methylome profiles using the Enzymatic Methyl-seq assay (EM-seq; NEBNext Enzymatic Methyl-seq; Illumina NovaSeq 6000 and NovaSeq X; 150bp paired-end reads; OICR Genomics Core)^8^.

**Figure 1.**
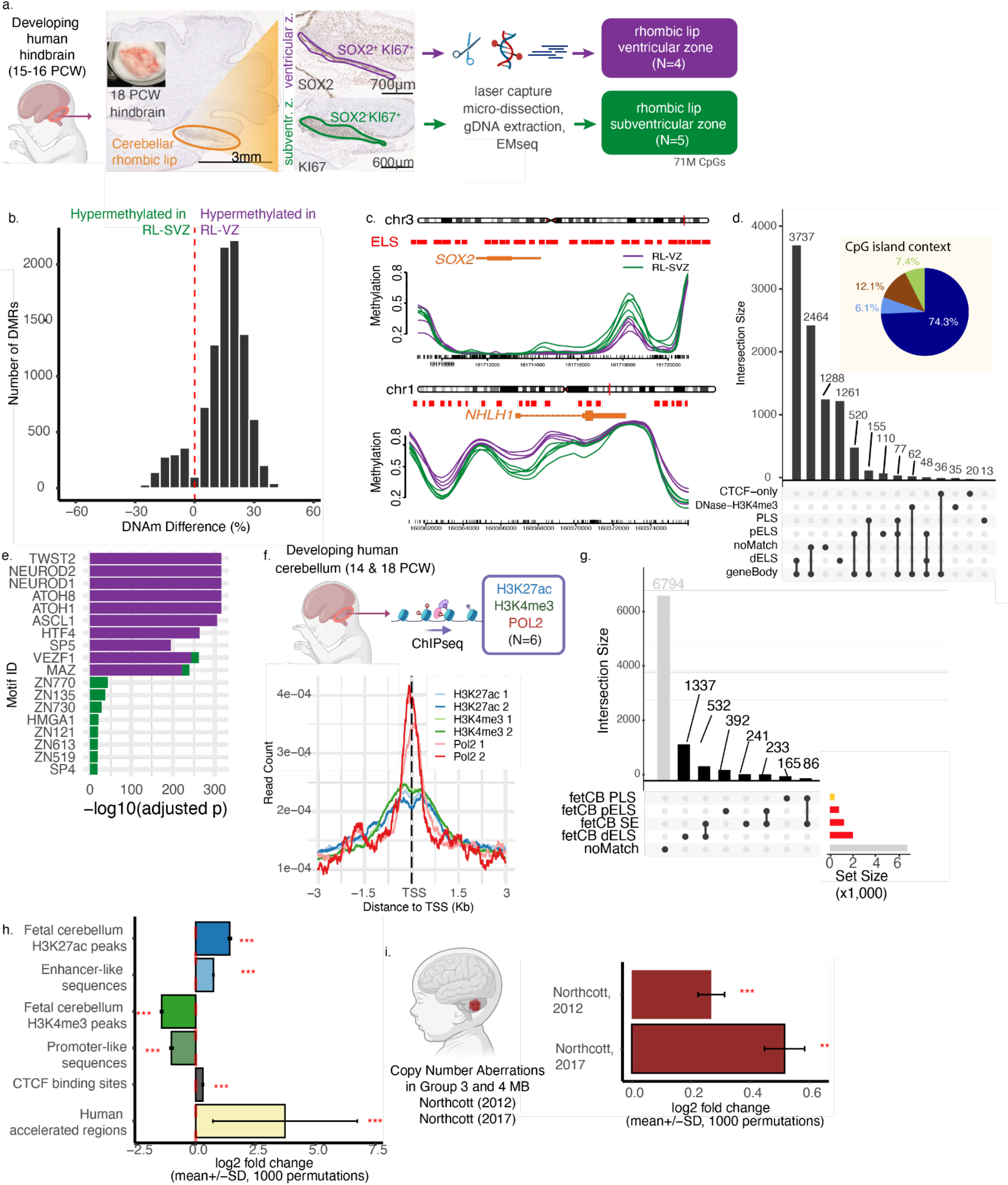
DNA methylome mapping of mid-gestation human rhombic lip reveals loci enriched in putative enhancers, human accelerated regions, and structural variants in medulloblastoma. a. Workflow to generate DNA methylomes from mid-gestation human rhombic lip ventricular zone (RL-VZ) and rhombic lip subventricular zone (RL-SVZ). b. Differentially methylated regions between RL-VZ and RL-SVZ (DSS, Q < 0.05). c. Sample-level smoothed methylation around *SOX2*, a representative locus hypermethylated in RL-SVZ (top), and *NHLH1*, hypermethylated in RL-VZ. (d) Overlap of DMRs with ENCODE consensus *cis* regulatory elements and gene bodies. Inset shows fraction of DMRs overlapping CpG islands (green), CpG shores (brown), shelfs (light blue), and open sea regions (deep blue). (e) Top ten most significant transcription factor binding site motifs for DMRs hypermethylated in RL-VZ (purple), and those for DMRs hypermethylated in RL-SVZ (green) (AME, Q < 0.05). (f) Schematic for generation of genome-wide chromatin immunoprecipitation data from human developing hindbrain (top) and average peak signal centered around transcription start sites (TSS). Each line indicates one technical replicate. (g) Overlap of DMRs with promoter-like and enhancer-like sequences as defined by human fetal cerebellar H3K4me3 and H3K27ac modifications and distance from TSS. (h) DMRs are enriched in enhancer-like regions, CTCF-binding sites, and human accelerated regions, and are depleted in promoter-like regions (permutation test, p < 10^-3^). (i) DMRs are enriched in copy number aberrations in Group 3 and 4 medulloblastoma (permutation test, p < 10^-3^). dELS: distal enhancer-like signatures; pELS: proximal enhancer-like signatures; PLS: promoter-like signatures; SE: super-enhancers.

### EM-seq data analysis

#### Processing

A standard bioinformatics pipeline was used to process the EMseq data; processing statistics are reported in Supplementary Table 2. Fastp 0.23.2^13^ was used to ascertain read quality and perform adapter trimming. Reads were aligned to the hg38 genome using bwa-meth 0.2.5^14^. Duplicates were filtered using Picard MarkDuplicates (https://broadinstitute.github.io/picard/). MethylDackel (https://github.com/dpryan79/MethylDackel) was used to compute Mbias plots. MethylDackel *extract --maxVariantFrac 0*.*2 --minOppositeDepth 5 --cytosine_report --OT 0,0,0,147 --OB: 3,0,5,0* was used to compute cytosine-level methylation reports while removing potential C > T variants. Methylation non-conversion rate was measured using the methylation level of the lambda phage genome. Samples had a mean non-conversion rate of 0.21% (range 0.34 - 0.64) and all samples were retained. All samples showed consistent patterns of genome-wide methylation with over 50% of CpGs having 90% methylation and 10% CpGs with <10% methylation (Supplementary Figure 2). Hierarchical clustering of CpG methylation in the region of SOX2 and EOMES largely separated the rhombic lip ventricular zone and subventricular zone samples, and no samples were excluded from downstream analysis (Supplementary Figure 1).

#### Differentially methylated regions

Differentially methylated regions (DMRs) were called for the CpG context using DSS^15^. Base-level hits were identified using DMLtest() with the following parameters: smoothing=TRUE, smoothing.span=500. Base-level statistics were merged into region-level hits using callDMR() with parameters *delta=0, p*.*threshold=1e-05, minlen=50, minCG=4, dis*.*merge=100, pct*.*sig=0*.*5*. We used AME for transcription factor binding site enrichment^16^. To test for enrichment of transcription factors active in the tissue of interest, we used the HOCOMOCO v12 CORE database as input after removing transcription factors expressed in less than 1% of rhombic lip ventricular or subventricular zone cells in a published snRNA-seq dataset^5^.

#### Region enrichment test

To compute statistical enrichment of DMRs in regions of interest, we used a permutation test. The fraction of DMR overlap with target regions was compared to those from a length- and GC-matched set of genomic intervals sampled from mappable regions of the genome (*gkmSVM::genNullSeqs()*). Regions with <90% overlap with Bismap-mappable regions of the genome^17^ were excluded. This process was repeated 1,000 times, using different random sets of genomic intervals. The fraction of times the null statistic was greater than or equal to that from the real data, was reported as the p-value.

#### Gene regulatory network inference

Predicted enhancer-gene links for H1-derived neural progenitor cells were downloaded from the companion website for Activity-by-Contact^9^ (see “External data and annotation sources” table below) and *liftOver* was used to map coordinates to hg38. DMRs were mapped to genes using these links. AME was used to infer transcription factor motifs overlapping a given DMR coordinate (ie., predicted enhancer) (union of *sequences*.*tsv* entries accompanying AME output for hypomethylated and hypermethylated DMRs, “true positive” sequences only). Where a DMR overlapped multiple TF binding sites, all were retained.

Predicted transcription factors were filtered for those differentially expressed between the rhombic lip ventricular zone and subventricular zone^3^ (For the network in Figure 2c: genes with FDR-adjusted p-value < 0.05; for the full table in Supplementary Table 9, genes with nominal p-value < 0.05).

**Figure 2.**
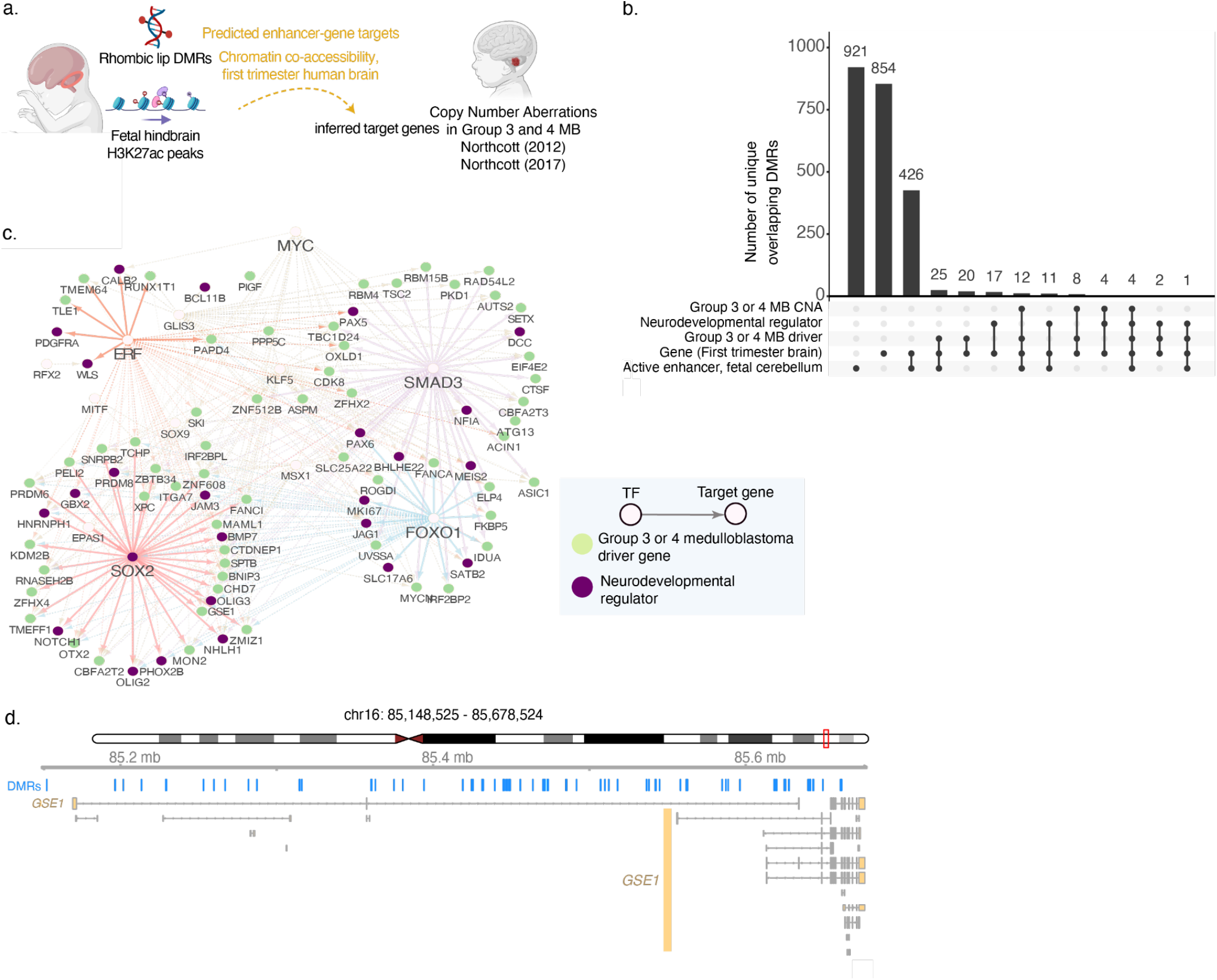
Annotation of differentially methylated regions with inferred target genes and medulloblastoma structural variants. a. Schematic for annotation. b. Overlap of differentially methylated regions (DMRs) with fetal cerebellum active enhancer marks, Overlap of putative rhombic lip candidate enhancers with fetal cerebellum active enhancer marks, cortical neural progenitor active enhancer marks and mutations in Group 3 and 4 medulloblastoma. Also annotated are regions where the nearest or predicted target gene is a known regulator of neurodevelopment or Group 3 and 4 medulloblastoma driver gene. c. Predicted gene regulatory network in the human rhombic lip where target genes are neurodevelopmental regulators (purple) or Group 3 and 4 medulloblastoma driver genes (yellow). Only links associated with genes of neurodevelopmental or cancer relevance are shown in this subnetwork (Supplementary Table 9). Selected targets of SOX2, FOXO1, SMAD3, and ERF are highlighted to display network substructure. d. Detailed view of rhombic lip DMRs at the GSE1 locus.

#### Association of promoter-level DNA methylation with gene expression

Processed RNAseq data for neuroanatomically-dissected rhombic lip ventricular zone and subventricular zone were downloaded from previously published work^3^. A standard bioinformatics pipeline and edgeR^18^ was used to call differentially expressed genes. Promoter level DNA methylation was compared to corresponding transcription levels in two different ways. First, for each gene, promoter-level DNA methylation was computed as the average methylation of the region 1000bp upstream and 150bp downstream of the TSS; locus-level methylation was averaged across samples within a neuroanatomical compartment and the methylation increase in rhombic lip ventricular zone (RL-VZ) was computed. This value was compared against gene expression level for genes significantly upregulated in the RL-VZ.

Separately, we limited analysis to just rhombic lip DMRs, and compared these to log2 fold-change of transcription in the RL-VZ.

### Human fetal cerebellum ChIP-seq data processing and peak calling

Acquisition of human tissue samples was approved by the Seattle Children’s Hospital Institutional Review Board. Two fresh frozen specimens from male fetal (14 and 18 PCW) human cerebellum were obtained from the Birth Defects Research Laboratory at the University of Washington with ethics board approval and maternal written consent obtained before specimen collection. Fresh frozen cerebellum (130-180 mg) was sent to Active Motif for ChIP-seq using the following antibodies: H3K4me3 (Active Motif, cat#39159), H3K27Ac (Active Motif, cat #39133) and RNA Pol II (Active Motif, cat # 91151).

#### Sequence Analysis

The 75-nt single-end (SE75) sequence reads generated by Illumina sequencing (using NextSeq 500) are mapped to the genome using the BWA algorithm^19^ (“bwa aln/samse” with default settings). Only reads that pass Illumina’s purity filter, align with no more than 2 mismatches, and map uniquely to the genome were used in the subsequent analysis. In addition, duplicate reads (“PCR duplicates”) were removed.

#### Determination of Fragment Density

Since the 5’-ends of the aligned reads (= “tags”) represent the end of ChIP/IP-fragments, the tags were extended *in silico* (using Active Motif software) at their 3’ ends to a length of 200 bp, which corresponds to the average fragment length in the size-selected library. To identify the density of fragments (extended tags) along the genome, the genome was divided into 32-nt bins and the number of fragments in each bin was determined.

#### Peak calling

Processing statistics are in Supplementary Table 10. Peaks were called using either MACS 2.1.0^20^ or SICER^21^ algorithms. MACS default cutoff was pvalue 1e-7 for narrow peaks and 1e-1 for broad peaks. SICER default cutoff was FDR 1e-10 with gap parameter of 600 bp. Peak filtering was performed by removing false ChIP-Seq peaks as defined within the ENCODE blacklist^22^.

Proximal H3K27ac peaks within 2 kbp upstream or downstream of transcription start sites that did not overlap with H3K4me3 peaks or distal H3K27ac peaks residing outside the +-2k bp window were annotated as enhancer-like elements. H3K4me3 peaks overlapping transcription start site +-200 bp windows were annotated as canonical promoter-like elements^23^. The elements were annotated for each sample and then combined for downstream analyses.

Superenhancers were called by the ActiveMotif pipeline. The identification of Super Enhancers uses a proprietary algorithm that gives a very similar result as the ROSE software. In a first step, MACS or SICER peaks generated by the standard ChIP-Seq analysis are merged (or “stitched together”) if their inner distance is equal or less than 12,500 bp. In the second step, the stitched peak regions with the strongest signals (top 5%) are identified as Super Enhancers. Super Enhancer intervals are ranked by their signal strength (= tag numbers in stitched peak regions). For each sample, the top-ranked Super Enhancer is thus at the top of the list.

### External data and annotation sources

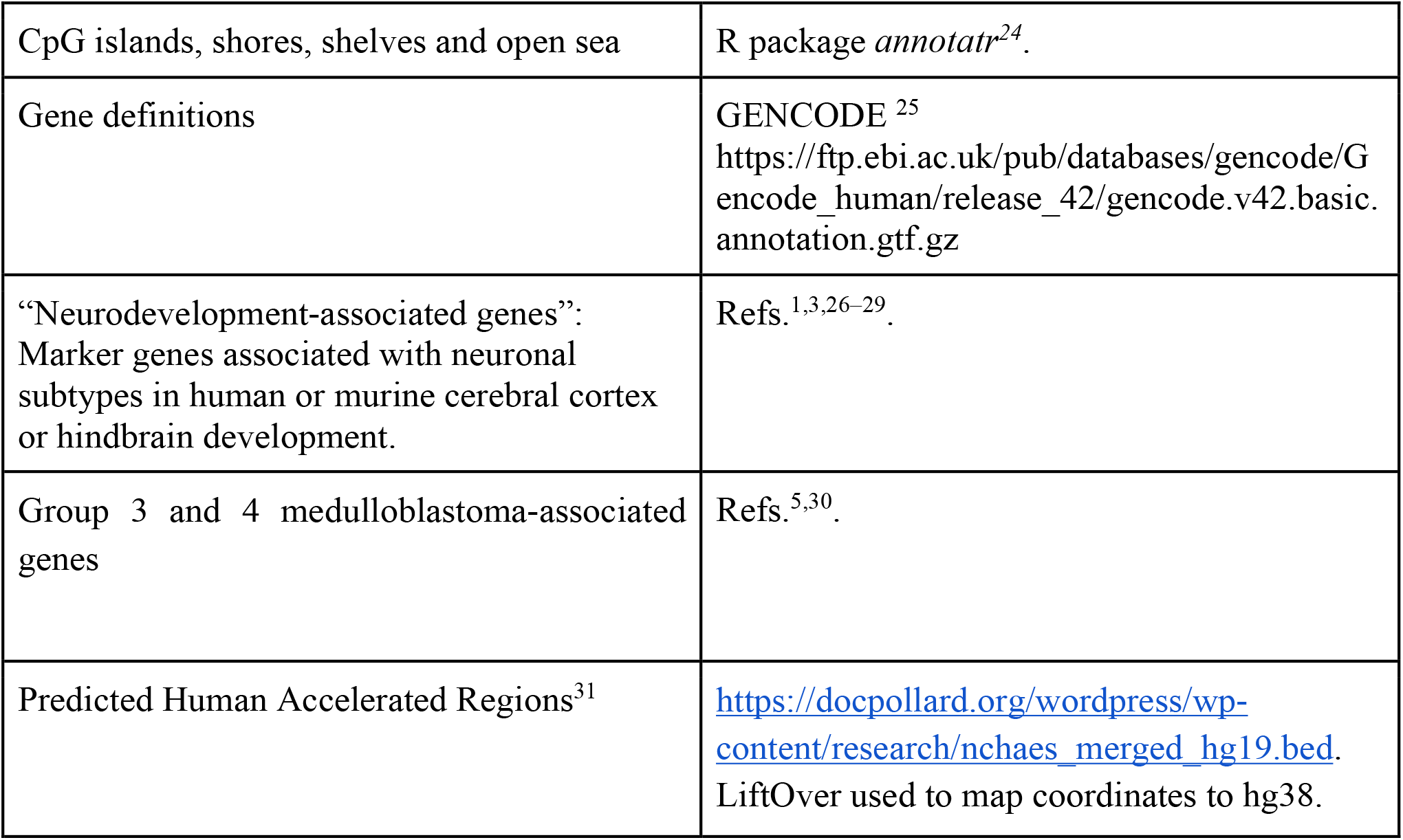
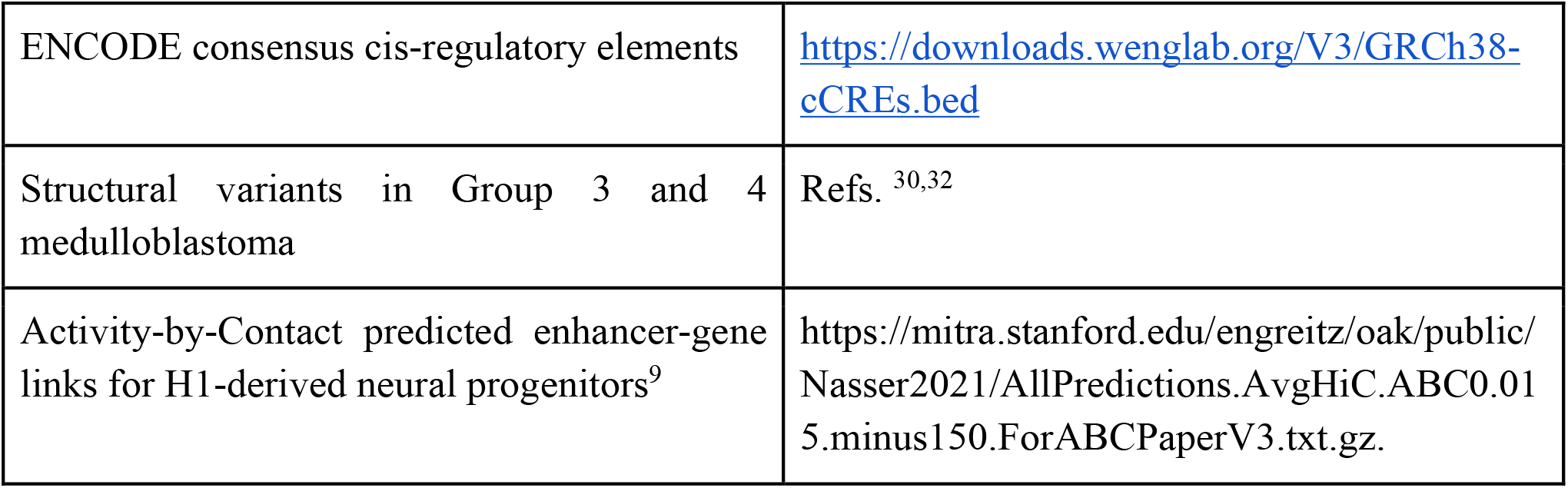

External data and annotation sources.
|  |  |
| --- | --- |
| CpG islands, shores, shelves and open sea | R package *annotatr*<sup>24</sup>. |
| Gene definitions | GENCODE <sup>25</sup><br>https://ftp.ebi.ac.uk/pub/databases/gencode/Gencode_human/release_42/gencode.v42.basic.annotation.gtf.gz |
| “Neurodevelopment-associated genes”:<br>Marker genes associated with neuronal subtypes in human or murine cerebral cortex or hindbrain development. | Refs. <sup>1,3,26–29</sup>. |
| Group 3 and 4 medulloblastoma-associated genes | Refs. <sup>5,30</sup>. |
| Predicted Human Accelerated Regions<sup>31</sup> | https://docpollard.org/wordpress/wp-content/research/nchaes_merged_hg19.bed.<br>LiftOver used to map coordinates to hg38. |
| ENCODE consensus cis-regulatory elements | https://downloads.wenglab.org/V3/GRCh38-cCREs.bed |
| Structural variants in Group 3 and 4 medulloblastoma | Refs. <sup>30,32</sup> |
| Activity-by-Contact predicted enhancer-gene links for H1-derived neural progenitors<sup>9</sup> | https://mitra.stanford.edu/engreitz/oak/public/Nasser2021/AllPredictions.AvgHiC.ABC0.015.minus150.ForABCPaperV3.txt.gz. |

## Results

### Regions showing epigenetic dynamics in the rhombic lip are enriched in putative active enhancers and structural variants of Group 3 and 4 medulloblastoma, and converge on transcription factors driving neurodifferentiation

We generated DNA methylomes for mid-gestation human rhombic lip ventricular and subventricular zones (**Fig. 1a**, N=9 samples from 5 males; 15-16 post-conception weeks; Supplementary Tables 1-2, Supplementary Figure 1-2) and looked for regions showing epigenetic dynamics over the course of rhombic lip differentiation. We identified 9,855 regions that showed significant differential CpG methylation (DMR) between the rhombic lip ventricular zone and rhombic lip subventricular zone (DSS^15^). The vast majority of DMRs were hypermethylated in the rhombic lip ventricular zone (88.5%), and were of an average length of 360bp (**Fig. 1b-c**; Supplementary Table 3; range of 51 - 4,122 bp length; mean = 46.3% hypomethylation). Figure 1c shows detailed views of sample-level methylation of DMRs in the genomic regions around *SOX2* and *NHLH1*, two genes known to regulate neurodifferentiation. Only 7.4% of DMRs overlapped CpG islands, and similar fractions overlapped CpG shores (12.1%) and shelves (6.1%); most DMRs were located outside these regions (74%; “open sea”)(**Fig. 1d**). We examined how DMRs were distributed among ENCODE consensus *cis* regulatory elements (CREs)(Supplementary Table 4)^23^. Over 80% of DMRs overlapped enhancer-like sequences, with the vast majority overlapping distal enhancer-like sequences (72.4%; 10.6% overlap proximal enhancer-like sequences). In contrast only 2.8% of DMRs overlapped promoters (**Fig. 1d**).

DMRs hypermethylated in the rhombic lip ventricular zone were enriched for binding sites for 339 transcription factors, with the top three being NEUROD1, NEUROD2, ATOH1 (**Fig. 1e**; AME, Q < 0.05; Supplementary Table 5). These transcription factors are known master regulators of rhombic lip cell identity and neuronal differentiation, with ATOH1 being an established regulator of glutamatergic cell identity in the rhombic lip ^1^. In contrast, DMRs hypermethylated in the rhombic lip subventricular zone are enriched for 159 transcription factor motifs, with the top 5 being ZNF770, ZNF135, ZNF730, HMGA1, and ZNF121 (AME Q <0.05; Supplementary Table 6).

We compared promoter-level DNA methylation change to change in gene expression, using a previously published transcriptome of the rhombic lip ventricular and subventricular zones^3^. Promoter-level DNA methylation demonstrates a weak negative correlation with gene expression (**Fig. S4a**, N=961 genes; Spearman’s rho = -0.075, p=0.022). Intriguingly, this correlation disappears if comparison is limited to the intersection of DMRs and differentially expressed genes (**Fig. S4b**, N=45 genes; Spearman’s rho = -0.15, p > 0.1).

To focus on the context of the developing cerebellum, we generated maps of histone modifications for active enhancers (H3K27ac), promoters (H3K4me3), and transcription (POL2) from the bulk fetal cerebellum (**Fig. 1f**, N=2 biological replicates, male, 14 and 18 post-conception weeks; Supplementary Table 10). We found that just under one-third of DMRs overlapped a putative enhancer or promoter region as defined by peaks of histone modifications in this tissue (31.1%, or 3,061 DMRs)(**Fig. 1g**). Most of these overlapped with peaks of active enhancer marks (24.4% of all DMRs), with only 8% overlapping putative promoters. Indeed, we found that DMRs were enriched in H3K27ac peaks in the fetal cerebellum, relative to length, GC- and mappability-matched sequences (**Fig. 1h**, p < 0.001). Importantly, DMRs were statistically depleted in H3K4me3 peaks in the human fetal cerebellum, a mark for active promoters (p < 0.001). We obtained the identical result using ENCODE CRE definitions of enhancers and promoters^23^ (p < 0.001). The fetal cerebellar H3K27ac maps revealed a set of 986 superenhancers (median length=26kb; range=2.8kb to 191kb; Supplementary Table 7). The nearest genes to these superenhancers included 28 genes previously linked to cerebellar neurodevelopment, including *NFIA, OTX2, EOMES, LMX1A*, and *ZIC1*. Notably, 37 of the superenhancers had known Group 3 and 4 medulloblastoma drivers as their nearest genes, including *CHD7, CBFA2T2, GSE1*, and of course, *OTX2*, and *ZIC1*. Notably, 11% of DMRs overlap regions predicted to contain superenhancers in the fetal cerebellum.

We next looked at DMR overlap with regions of accelerated evolution in the human genome, and with DNA copy number aberrations in Group 3 and 4 medulloblastoma. We found that 25 DMRs overlap human accelerated regions (Supplementary Table 8)^31^. These include regions with nearest protein-coding genes such as *BCAS3, RFX3, SOBP*, and *PEX14*, mutations in which have been linked to intellectual disability, neurological deficits, attention deficit hyperactivity disorder, or autism spectrum disorders^33–36^. To examine enrichment of DMRs in copy number aberrations, we considered copy number aberrations (CNA) from two independent publications and called using two different genomic platforms^30,32^. We found that DMRs were enriched in both sets of CNA peaks tested (p < 0.001 in both instances). Close to one-quarter of the DMRs overlap known amplifications and deletions in Group 3 and 4 medulloblastoma (N=2,135 DMRs, or 22%). Notably, 61 DMRs overlap a 530kb window containing genetic suppressor element 1 (*GSE1*), a region also predicted to contain multiple superenhancers. GSE1 encodes a coiled-coil domain protein, and is associated with the CoREST transcription repressor complex in mouse placental development^37^. Frameshift loss-of-function mutations in GSE1 are a known driver in the *PTCH1*-mutated Sonic Hedgehog subtype of medulloblastoma^38,39^, suggesting a role for this protein in development and oncogenesis. The complete list of DMRs annotated with overlapping target genes and tumour CNAs is provided in Supplementary Table 9. Rhombic lip DMRs located in copy number aberrations may point to cis-regulatory DNA elements that may be amplified or deleted in medulloblastoma, resulting in a functional change in gene regulation, and in driving the cancer.

Collectively, our data provide evidence that regions showing DNA methylation dynamics within the mid-gestation human rhombic lip are enriched for enhancers, regions showing accelerated evolution in humans, and large-scale genomic aberrations in Group 3 and 4 medulloblastoma. We next used these data as a starting point to narrow down individual candidate regions as high-confidence enhancer predictions.

### Predicted cerebellar rhombic lip gene regulatory network identifies new transcription factor links to developmental genes and medulloblastoma driver genes

We next sought to analyze the DMRs to nominate a list of predicted enhancers and their target genes. For this purpose, we integrated the rhombic lip specific DMRs with the fetal cerebellar histone maps generated above, with previously predicted enhancer-gene pairs from multimodal epigenomic data^9,10^, and with chromatin co-accessibility data from the first trimester brain^12^ (**Fig. 2**, Supplementary Table 9). 14% of DMRs overlap active enhancer marks, as defined by H3K27ac peaks in the fetal cerebellum (**Fig. 2a-b**, N=1,400 DMRs). Separately, 14% of DMRs are predicted to regulate the expression of 4,604 genes (N=1,384 DMRs)^9^.

As our interest is in identifying non-coding DNA elements that impact the regulation of genes involved in neurodifferentiation and medulloblastoma oncogenesis, we predicted transcription factor-enhancer-target gene links (triplet links) using our set of rhombic lip DMRs^9,16^ (Supplementary Table 9; subset shown in **Fig. 2c**). We limited our transcription factor set to genes previously shown to be differentially expressed between the rhombic lip ventricular zone and subventricular zone^3^. We identified a total of 81,844 triplet links, covering over 98% of DMRs (9,741 DMRs). The links are associated with 41 transcription factors and include 4,610 predicted target genes. We identified 29 DMR target genes previously associated with neurodevelopmental regulation, including *SOX2, WLS, PAX6*, and *MKI67* (**Supplementary Table 9**). Our network includes previously known gene regulatory connections such as the regulation of *Notch1* and *Otx2* by Sox2^40,41^, and regulation of *PAX6* by SMAD3^42^. including the regulation of DMRs were also predicted to regulate the RUNX1 transcriptional co-repressors *CBFA2T2* and *CBFA2T3*, which have been shown demarcate the rhombic lip subventricular zone. *CBFA2T2* and *CBFA2T3* are recurrently mutated in Group 4 medulloblastoma, a tumour hypothesized to originate in the rhombic lip subventricular zone. Our triplet links also include 60 genes known to be recurrently mutated or overexpressed in medulloblastoma, including *OTX2, MYCN*, the Fanconi anemia proteins *FANCA* and *FANCI*, and others (**Fig. 2c, Supplementary Table 9**).

## Discussion

We have mapped, to our knowledge, the first DNA methylomes of the neuroanatomically-dissected human cerebellar rhombic lip, the progenitor niche which goes on to generate over 80% of the neurons in the adult human brain, and the dysregulation of which is hypothesized to result in structural cerebellar birth defects and medulloblastoma. The advantage of our approach is that, unlike the cell type inference required from single-cell genomics datasets, here we directly sample neuroanatomical compartments of the rhombic lip ventricular and subventricular zones. From these data, we infer gene regulatory networks that include 41 transcription factors active in the rhombic lip, and target genes that mediate neurodevelopment and are drivers of Group 3 and 4 medulloblastoma. By integrating methylome maps with novel histone maps of the human fetal cerebellum that mark active enhancers and promoters, we predicted nearly 1,400 enhancers that regulate over 4,600 genes. These include enhancers of regulators of genes that drive glutamatergic neuronal identity (e.g., PAX6, WLS), and known drivers of medulloblastoma (OTX2, MYCN, CBFA2T2). These data provide a rich starting point for functional studies to dissect the gene regulatory impact of non-coding variation observed in genomes of medulloblastoma tumours and patients with structural birth defects of the cerebellum.

The presence of a weak negative correlation between DNA methylation at the promoter and change in transcription levels has been noted in other cellular contexts, and may be due to multiple factors^43^, such as tissue and cell type. Another possible confounding source is due to variation in the size of gene-specific promoters as well as the nature of the DNA modification measured. Enzymatic Methyl-seq cannot distinguish between 5-methylcytosine and 5-hydroxymethylcytosine, which show opposite correlations with transcription levels^44^. Moreover, rhombic lip DMRs appear to be enriched in putative enhancers, and depleted in promoters. Functional studies are needed to investigate the impact of altering putative transcription factor binding sites in these loci and on gene expression in the rhombic lip. Collectively, our results point to an epigenetic modulation of genes between the rhombic lip ventricular and subventricular zone, rather than a binary switch on and off by altered methylation at the promoters.

One limitation of the current study is that we used only male samples to generate DNA methylomes. Future studies will need to include female samples to ascertain if sex-specific epigenomic differences can explain the bias in incidence of some subtypes of medulloblastoma in males^45^. Another limitation of the current study is that we integrate DMRs from the neuroanatomically distinct and small glutamatergic rhombic lip, with histone marks from the whole fetal cerebellum. A large proportion of the latter includes GABAergic cells such as the Purkinje cell lineage, glutamatergic neurons in later states of differentiation (granule cells), and non-neuronal cell populations^1,26^. As technologies for low-input histone mapping become more widely used, our predictions can be refined using annotations that are more specific to rhombic lip cells.

Functional validation of disease variants predicted to impact gene expression will be necessary for eventual use of this information in genomic diagnostics^46,47^.

## Acknowledgements

We acknowledge the NIH NeuroBioBank and the University of Maryland Brain and Tumour Bank for the fetal tissue samples from which the DNA methylomes were generated. We thank Drs. Kathleen Millen and Parthiv Haldipur for guidance in the neuroanatomical isolation of the rhombic lip ventricular zone and subventricular zone, and Dr. Kathleen Millen for feedback on this manuscript. This work was funded by an Ontario Institute of Cancer Research Investigator Award, Cancer Research Society Operating Award (# 1058075), National Sciences and Engineering Research Council Discovery Grant (DGECR-2022-00236), and Canadian Institutes of Health Research Priority Award (PDI #185647) to S.P. We acknowledge the Birth Defects Research Laboratory (BDRL) at the University of Washington, including Ian A. Glass, Kimberly A. Aldinger, Dan Doherty, Ian Phelps, Jennifer Dempsey, Yasmeen Otaibi, and Eric Y. So with funding provided by NIH under NICHD R24HD000836 (I.A.G.). This study was conducted with the support of the Ontario Institute for Cancer Research’s Genomics Program (genomics.oicr.on.ca) through funding provided by the Government of Ontario. BioRender was used to generate some of the figures in this work.

## Data and Code Availability

Upon publication, fetal hindbrain EMseq raw data, aligned reads and methylation counts will be deposited in the Gene Expression Omnibus. Fetal cerebellum H3K7ac, H3K4me and POL2 ChIPseq data will be deposited in the Gene Expression Omnibus. Called DMRs have been made available in supplementary data associated with this manuscript.

Software to reproduce analysis in this manuscript is available at https://github.com/RealPaiLab/RhombicLip_Epigenome and will be made available under a Creative Commons Attributions License upon publication.

## Supplementary Figures

**Supplementary Figure 1.**
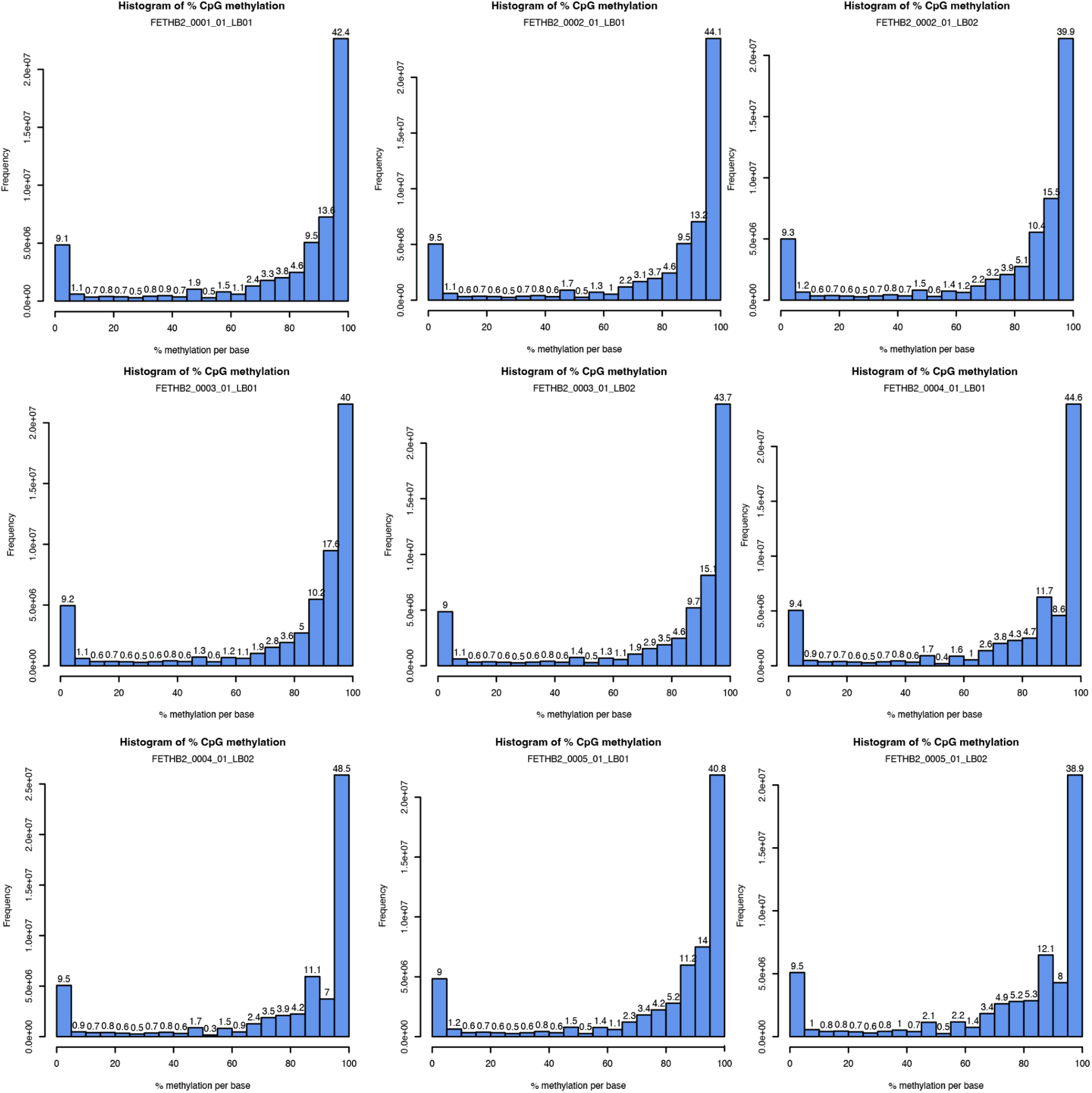
Genome-wide distribution of base-level percent DNA methylation. Each panel shows data for an individual DNA methylome.

**Supplementary Figure 2.**
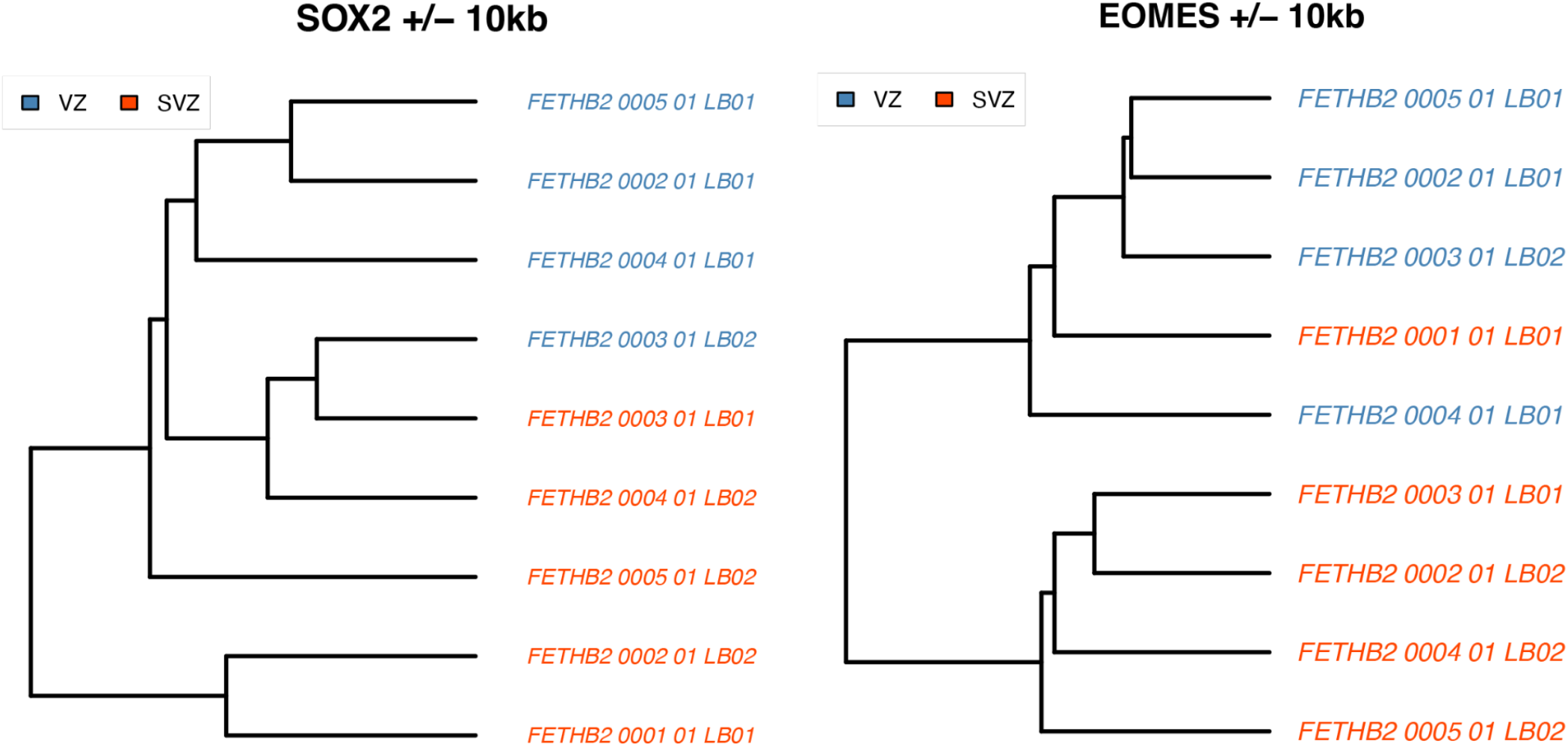
Hierarchical clustering of DNA methylomes based on CpG methylation of (a) the SOX2 gene region (b) EOMES gene region.

**Supplementary Figure 3.**
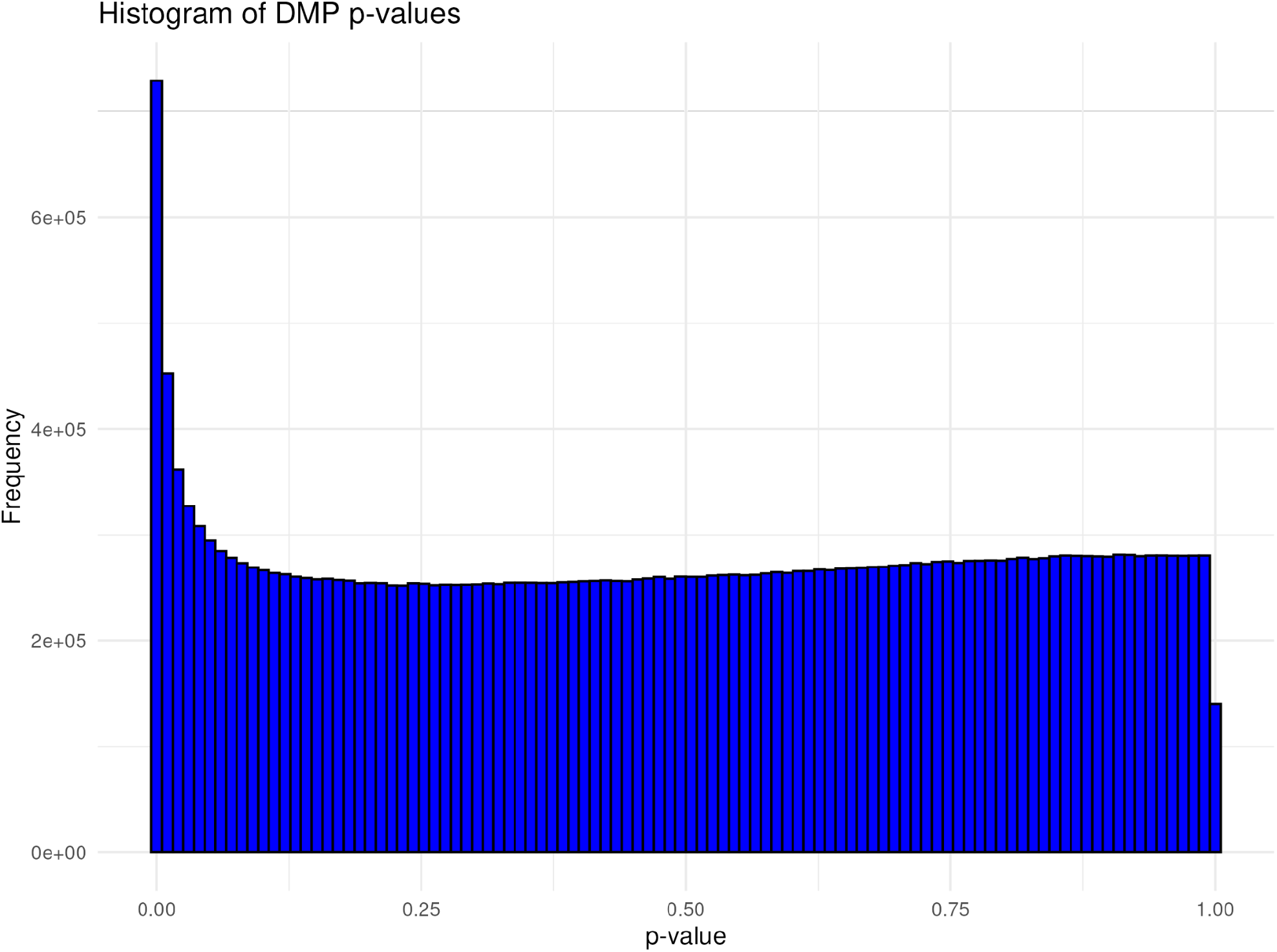
Distribution of nominal p-values of base-level differentially methylated CpGs between rhombic lip ventricular zone and subventricular zone.

**Supplementary Figure 4.**
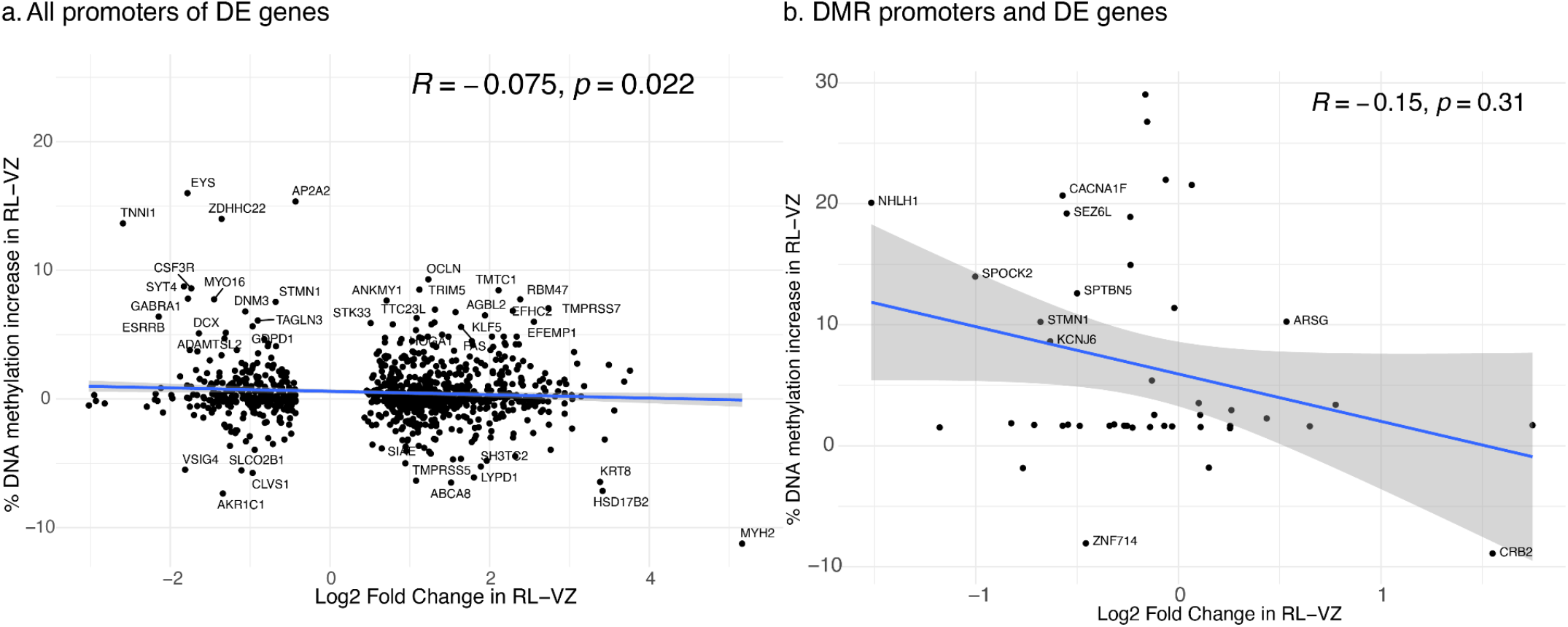
Promoter-level DNA methylation and gene expression. a. Promoter-level DNA methylation increase and transcription increase of the corresponding gene; analysis limited to significantly differentially expressed genes (N=961 genes). Spearman correlation and corresponding p-value from a t-test are shown. b. Promoter-level DNA methylation increase and transcription increase are limited to genes where the promoter region overlaps a DMR and which is differentially expressed in the rhombic lip (N=45 genes). Spearman correlation and corresponding p-value from a t-test are shown.

## References

1. Leto, K. et al. Consensus paper: Cerebellar development. Cerebellum 15, 789–828 (2016).

2. Haldipur, P. et al. Evidence of disrupted rhombic lip development in the pathogenesis of Dandy-Walker malformation. Acta Neuropathol. 142, 761–776 (2021).

3. Aldinger, K. A. et al. Spatial and cell type transcriptional landscape of human cerebellar development. Nat. Neurosci. (2021) doi:10.1038/s41593-021-00872-y.

4. Williams, R. W. & Herrup, K. The control of neuron number. Annu. Rev. Neurosci. 11, 423–453 (1988).

5. Hendrikse, L. D. et al. Failure of human rhombic lip differentiation underlies medulloblastoma formation. Nature 609, 1021–1028 (2022).

6. Haldipur, P. et al. Spatiotemporal expansion of primary progenitor zones in the developing human cerebellum. Science 366, 454–460 (2019).

7. Smith, K. S. et al. Unified rhombic lip origins of group 3 and group 4 medulloblastoma. Nature 609, 1012–1020 (2022).

8. Vaisvila, R. et al. Enzymatic methyl sequencing detects DNA methylation at single-base resolution from picograms of DNA. Genome Res. 31, 1280–1289 (2021).

9. Nasser, J. et al. Genome-wide enhancer maps link risk variants to disease genes. Nature 593, 238– 243 (2021).

10. Fulco, C. P. et al. Activity-by-contact model of enhancer-promoter regulation from thousands of CRISPR perturbations. Nat. Genet. 51, 1664–1669 (2019).

11. Roadmap Epigenomics Consortium et al. Integrative analysis of 111 reference human epigenomes. Nature 518, 317–330 (2015).

12. Mannens, C. C. A. et al. Chromatin accessibility during human first-trimester neurodevelopment. Nature 1–8 (2024).

13. Chen, S., Zhou, Y., Chen, Y. & Gu, J. fastp: an ultra-fast all-in-one FASTQ preprocessor. Bioinformatics 34, i884–i890 (2018).

14. Pedersen, B. S., Eyring, K., De, S., Yang, I. V. & Schwartz, D. A. Fast and accurate alignment of long bisulfite-seq reads. arXiv [q-bio.GN] (2014).

15. Park, Y. & Wu, H. Differential methylation analysis for BS-seq data under general experimental design. Bioinformatics 32, 1446–1453 (2016).

16. Machanick, P. & Bailey, T. L. MEME-ChIP: motif analysis of large DNA datasets. Bioinformatics 27, 1696–1697 (2011).

17. Karimzadeh, M., Ernst, C., Kundaje, A. & Hoffman, M. M. Umap and Bismap: quantifying genome and methylome mappability. Nucleic Acids Res. 46, e120 (2018).

18. Chen, Y., Chen, L., Lun, A. T. L., Baldoni, P. L. & Smyth, G. K. edgeR v4: powerful differential analysis of sequencing data with expanded functionality and improved support for small counts and larger datasets. bioRxiv (2024) doi:10.1101/2024.01.21.576131.

19. Li, H. & Durbin, R. Fast and accurate short read alignment with Burrows-Wheeler transform. Bioinformatics 25, 1754–1760 (2009).

20. Feng, J., Liu, T. & Zhang, Y. Using MACS to identify peaks from ChIP-Seq data. Curr. Protoc. Bioinformatics Chapter 2, 2.14.1–2.14.14 (2011).

21. Zang, C. et al. A clustering approach for identification of enriched domains from histone modification ChIP-Seq data. Bioinformatics 25, 1952–1958 (2009).

22. ENCODE Project Consortium. An integrated encyclopedia of DNA elements in the human genome. Nature 489, 57–74 (2012).

23. ENCODE Project Consortium et al. Expanded encyclopaedias of DNA elements in the human and mouse genomes. Nature 583, 699–710 (2020).

24. Cavalcante, R. G. & Sartor, M. A. Annotatr: Genomic regions in context. Bioinformatics 33, 2381– 2383 (2017).

25. Frankish, A. et al. GENCODE: reference annotation for the human and mouse genomes in 2023. Nucleic Acids Res. 51, D942–D949 (2023).

26. Sepp, M. et al. Cellular development and evolution of the mammalian cerebellum. Nature 625, 788– 796 (2024).

27. Bhaduri, A. et al. An atlas of cortical arealization identifies dynamic molecular signatures. Nature 598, 200–204 (2021).

28. Vladoiu, M. C. et al. Childhood cerebellar tumours mirror conserved fetal transcriptional programs. Nature 572, 67–73 (2019).

29. Carter, R. A. et al. A Single-Cell Transcriptional Atlas of the Developing Murine Cerebellum. Curr. Biol. 28, 2910–2920.e2 (2018).

30. Northcott, P. A. et al. The whole-genome landscape of medulloblastoma subtypes. Nature 547, 311– 317 (2017).

31. Capra, J. A., Erwin, G. D., McKinsey, G., Rubenstein, J. L. R. & Pollard, K. S. Many human accelerated regions are developmental enhancers. Philos. Trans. R. Soc. Lond. B Biol. Sci. 368, 20130025 (2013).

32. Northcott, P. A. et al. Subgroup-specific structural variation across 1,000 medulloblastoma genomes. Nature 488, 49–56 (2012).

33. Harris, H. K. et al. Disruption of RFX family transcription factors causes autism, attention-deficit/hyperactivity disorder, intellectual disability, and dysregulated behavior. Genet. Med. 23, 1028–1040 (2021).

34. Birk, E. et al. SOBP is mutated in syndromic and nonsyndromic intellectual disability and is highly expressed in the brain limbic system. Am. J. Hum. Genet. 87, 694–700 (2010).

35. Hengel, H. et al. Bi-allelic loss-of-function variants in BCAS3 cause a syndromic neurodevelopmental disorder. Am. J. Hum. Genet. 108, 1069–1082 (2021).

36. Waterham, H. R. et al. Autosomal dominant Zellweger spectrum disorder caused by de novo variants in PEX14 gene. Genet. Med. 25, 100944 (2023).

37. Hiver, S. et al. Gse1, a component of the CoREST complex, is required for placenta development in the mouse. Dev. Biol. 498, 97–105 (2023).

38. Skowron, P. et al. The transcriptional landscape of Shh medulloblastoma. Nat. Commun. 12, 1749 (2021).

39. Huang, M. et al. Engineering genetic predisposition in human neuroepithelial stem cells recapitulates medulloblastoma tumorigenesis. Cell Stem Cell 25, 433–446.e7 (2019).

40. Taranova, O. V. et al. SOX2 is a dose-dependent regulator of retinal neural progenitor competence. Genes Dev. 20, 1187–1202 (2006).

41. Chan, C. S. Y. et al. Cell type- and stage-specific expression of Otx2 is regulated by multiple transcription factors and cis-regulatory modules in the retina. Development 147, dev187922 (2020).

42. Qian, Z. et al. Investigating the mechanism by which SMAD3 induces PAX6 transcription to promote the development of non-small cell lung cancer. Respir. Res. 19, 262 (2018).

43. Anastasiadi, D., Esteve-Codina, A. & Piferrer, F. Consistent inverse correlation between DNA methylation of the first intron and gene expression across tissues and species. Epigenetics Chromatin 11, 37 (2018).

44. Khare, T. et al. 5-hmC in the brain is abundant in synaptic genes and shows differences at the exonintron boundary. Nat. Struct. Mol. Biol. 19, 1037–1043 (2012).

45. Cavalli, F. M. G. et al. Intertumoral Heterogeneity within Medulloblastoma Subgroups. Cancer Cell 31, 737–754.e6 (2017).

46. Ellingford, J. M. et al. Recommendations for clinical interpretation of variants found in non-coding regions of the genome. Genome Med. 14, 73 (2022).

47. Klein, J. C. et al. A systematic evaluation of the design and context dependencies of massively parallel reporter assays. Nat. Methods 17, 1083–1091 (2020).

